# The upper respiratory tract microbiome of Australian Aboriginal and Torres Strait Islander children in ear and nose health and disease

**DOI:** 10.1101/2021.05.13.444113

**Authors:** Andrea Coleman, Julian Zaugg, Amanda Wood, Kyra Cottrell, Eva Grahn Håkansson, Jasmyn Adams, Matthew Brown, Anders Cervin, Seweryn Bialasiewicz

**Affiliations:** The University of Queensland Centre for Clinical Research; Herston (4001), Australia; Townsville University Hospital, Townsville (4814), Australia; Australian Centre for Ecogenomics, The University of Queensland, St Lucia (4067), Australia; Queensland Health Deadly Ears Program, Brisbane (4001), Australia; Clinical Microbiology, Umeå University, Umeå (901 87), Sweden; The Royal Brisbane and Women’s Hospital, Brisbane (4001), Australia; Queensland Paediatric Infectious Diseases Laboratory, Queensland Children’s Hospital, South Brisbane (4001), Australia

## Abstract

**Objective:** To examine the nasal microbiota in relation to otitis status and nose health in Indigenous Australian children.

**Methods:** Children aged 2-7 years were recruited from two northern Australian (Queensland) communities. Clinical histories were obtained through parent interview and review of the medical record. Nasal cavity swabs were obtained, and the child’s ears, nose and throat were examined. DNA was extracted and analysed by 16S rRNA amplicon next generation sequencing of the V3/V4 region in combination with previously generated culture data.

**Results:** 103 children were recruited (mean 4.7 years), 17 (16.8%) were ‘healthy’, i.e. normal examination and no history of otitis media (OM). Nasal microbiota differed significantly in relation to otitis status and nose health. Children with historical OM had higher relative abundance of *Moraxella* compared to healthy children, despite both having healthy ears at the time of swabbing. Children with healthy noses had higher relative abundance of *S. aureus* compared to those with rhinorrhoea. *Dolosigranulum* was correlated to *Corynebacterium* in healthy children. *Haemophilus* and *Streptococcus* correlated across phenotypes. *Ornithobacterium* was absent/low relative abundance in healthy children and clustered around otopathogens. It correlated with *Helcococcus* and *Dichelobacter*.

**Conclusions:** *Dolosigranulum* and *Corynebacterium* form a synergism that promotes URT/ear health in Indigenous Australian children. *Ornithobacterium* likely represents *Candidatus Ornithobacterium hominis* and in this population is correlated with a novel bacterium which appears to be related to poor upper respiratory tract/ear health.

**Importance:** Recurring and chronic infections of the ear (otitis media) are disproportionately prevalent in disadvantaged communities across the globe, and in particular, within Indigenous communities. Despite numerous intervention strategies, otitis media persists as a major health issue and is the leading cause of preventable hearing loss. In disadvantaged communities, this hearing loss is associated with negative educational and social development outcomes, and consequently, poorer employment prospects and increased contact with the justice system in adulthood. Thus, a better understanding of the microbial ecology is needed in order to identify new targets to treat, as well as prevent the infections. This study used a powerful combination of 16S rRNA sequencing and extended culturomics to show that *Dolosigranulum pigrum*, a bacterium previously identified as a candidate protective species, may require co-colonisation with *Corynebacterium pseudodiptheriticum* in order to prevent otitis media. Additionally, emerging and potentially novel pathogens and bacteria were identified.

## Introduction

Otitis media (OM), an inflammation/infection of the middle ear, is a common paediatric condition[1]. However, in many indigenous populations globally there is a disproportionately high OM-associated burden, impacting negatively on schooling and employment outcomes[1, 2]. Previous microbiological studies relating to OM in indigenous populations have been largely limited to the main otopathogens (*Streptococcus pneumoniae, Haemophilus influenzae*, and *Moraxella catarrhalis*) using culture-dependent methods and seldom included healthy indigenous control children[3]. One study used 16S ribosomal RNA (rRNA) next generation sequencing (NGS) to explore the middle ear effusion and nasopharyngeal/adenoid microbiota in relation to OM with effusion (OME) in 11 Aboriginal and/or Torres Strait Islander (referred to herein as Indigenous Australian) children[4], which confirmed the association of otopathogen-containing genera and OME.

We have previously used culturomics and species-specific quantitative PCR (qPCR) to explore the nasal microbiota in relation to ear health and OM in 103 Indigenous Australian children[5]. We found that children with historical or current OM/upper respiratory tract (URT) infection (URTI) had high otopathogen loads and higher detection of rhinovirus[5]. In contrast, *Corynebacterium pseudodiphtheriticum* and *Dolosigranulum pigrum* were associated with URT/ear health[5]. However, culture-based analyses can be insensitive to microbial population structure and fastidious or unculturable organisms, such as the recently described, *Candidatus Ornithobacterium hominis*[6, 7]. To address this limitation, 16S rRNA NGS, supplemented with the existing culturomics data, was used to investigate the broader bacterial microbiome and how it relates to ear and nose health and disease in Indigenous Australian children.

## Materials and Methods

Additional detail of the materials and methods can be found in the supplementary material.

### Population and sample collection

Indigenous Australian children aged 2–7 years old were recruited prospectively from one rural and one remote northern Queensland communities in Australia through October 2015– November 2017. Children whom received antibiotics within three weeks of sample collection were excluded[5]. The study was approved by the Far North Queensland Human Research Ethics Committee (HREC/15/QCH/10-594).

A detailed description of the cohort, sampling and clinical data collection has been previously documented[5]. Briefly, demographic details and ear health history were collected for eligible children from parent interview and the child’s medical record. Children underwent ear (otoscopy), nose and throat (ENT) examination. Ear status at time of swabbing was classified according to the most affected ear. Intra-nasal mucosal swabs (dry FLOQSwabs, Copan Diagnostics, USA) used for molecular analysis were collected in parallel with Rayon swab (Transystem™ Minitip, Copan Diagnostics, USA) for culturomics[5]. All swabs were kept at 4°C from time of collection until arrival at the laboratory 24–48 hours later. Molecular swabs were then stored at -80°C.

### DNA extraction and quality assurance

DNA was extracted via mechanical bead beating and tissue lysis, followed by automated MagNa Pure (Roche Diagnostics, Australia), as previously described[5]. Four clean negative control swabs were processed in parallel with the sample swabs. The quality of nasal sampling was assessed using a real-time PCR targeting the endogenous retrovirus-3 (ERV3) marker for human DNA[8]. Swabs that amplified with cycle thresholds ≤38 were considered to have adequate nasal epithelial cell content, and by extension, be of good collection quality. Swabs producing cycle thresholds >38 were excluded from further analysis.

#### 16S Sequencing

All sample and negative control DNA extracts underwent 16S rRNA gene amplification using the 341F;806R primer set, followed by secondary indexing PCR. The equimolar library pool was then sequenced on an Illumina MiSeq (San Diego, CA, USA) with a V3, 600 cycle kit (2 x 300 base pairs paired-end).

#### Sequence data processing

Primer sequences were removed from de-multiplexed reads using cutadapt (ver. 2.6). Using QIIME2 (ver. 2019.10.0), reads were filtered, dereplicated and chimeras removed by DADA2. Taxonomy was assigned to the resulting amplicon sequence variants (ASV) by aligning each (classify-consensus-blast) against the non-redundant SILVA database (release 138).

#### Data availability

Amplicon sequencing data has been deposited in NCBI’s Short Read Archive under BioProject number PRJNA684919 (https://www.ncbi.nlm.nih.gov/sra/PRJNA684919) with accession numbers SRR13264782-SRR13264885.

#### Data analysis and statistics

Amplicon data analyses were performed in R (ver. 4.0.2). ASVs that were not bacterial, fungal or archaeal in origin, classified at below the phylum level, or that were classified as chloroplast or mitochondria, were discarded. Putative contaminants were identified using decontam (ver. 1.8.0) and microdecon (ver. 1.0.2) and removed. ASVs with a relative abundance <0.05% were also removed, with those samples with less than 4,000 reads remaining then discarded. Sample depth was limited to a maximum of 50,000 reads by using the rrarefy function in vegan (ver. 2.5-6).

Vegan was used to perform principal-component analysis (PCA), permutational multivariate analysis of variance (PERMANOVA) and analysis of multivariate homogeneity (PERMDISP) on centred log-ratio (clr) transformed ASV counts collapsed to the genus level. Differentially abundant ASVs and genera were identified using DESeq2 (ver. 1.28.1). Alpha diversity metrics Chao1, Shannon, and Simpson were calculated using phyloseq (ver. 1.32.0) on samples rarefied to 10,000 reads. Significant differences in alpha diversity distributions were determined through either Mann-Whitney U tests or Kruskal-Wallis and Dunn’s multiple comparisons tests, corrected for multiple testing using the Benjamini and Hochberg method. FastSpar (ver. 0.0.10) was used for correlation analysis of genera.

### Culturomic analysis

Culture-based swabs were processed using an expanded agar protocol under aerobic and anaerobic conditions with Vitek MS MALDI-TOF (bioMérieux) isolate identification as previously described[5]. Agreement between culture and 16S sequencing was assessed using Cohen’s Kappa.

## Results

In total, 103 children were recruited; two children refused swabbing, resulting in 101 swabs for analysis. All swabs met quality assurance criteria on ERV3 testing. Raw sample 16S read depth ranged from 149–262,880 (median 119,693), with quality, contamination and non-specific filtering resulting in the remaining read depth ranging from 0–163,794 (median 66,929). Fourteen samples were subsequently excluded as they did not pass quality control. The agreement between culturomics and 16S sequencing was 59.2%, Cohen’s kappa 0.08. The low level of agreement was predominately due to high sensitivity of detection by 16S sequencing, detecting on average of 14.3 (range 1–73) more genera per sample compared to culture.

### Nasal Microbiota in relation to ear health

Only 17 children (16.8%) had no history of OM and normal ENT examinations at the time of swabbing (Never OM), 7 (6.9%) had a perforated TM, 18 (17.8%) had a middle ear effusion (Effusion), 4 (4.0%) had AOM, and 55 had a past history of OM, but normal TM at the time of swabbing (HxOM) (54.5%) (Table 1). Due to low numbers, AOM samples were excluded from further analyses. There was a significant difference in the nasal microbiota in relation to otitis status (PERMANOVA F = 2.101, *p* = 0.0027), although with dispersion differences (PERMDISP F = 3.341, *p* = 0.0244). Within children that had healthy TMs at the time of sampling, HxOM had higher mean abundance of *Moraxella* compared to Never OM (31.22% vs 20.22%, p<0.05) (Figure 1, Supplementary Table 1). The relative abundance of nine *Dolosigranulum* ASVs differed significantly in relation to otitis status; ASVs 588 and 2067 were more abundant in children with normal TMs, while ASVs including 1030, 1069 and 1528 were more abundant in children with OM (Supplementary Table 2) The relative abundance of *Dolosigranulum* was positively correlated with *Corynbacterium* in Never OM and both *Corynbacterium* and *Moraxella* in HxOM; there was no significant correlation between *Dolosigranulum* and the other main otopathogen-containing genera (Supplementary Figure 1). Children with Effusion had higher mean relative abundance of *Ornithobacterium* (34.1%) compared to Never OM (28.4%); although non-significant according to DESeq, was significant according to Dunn’s test (adjusted *p* = 0.018, KrusW *p* = 0.021) (Supplementary Figure 2).

**Table 1:**
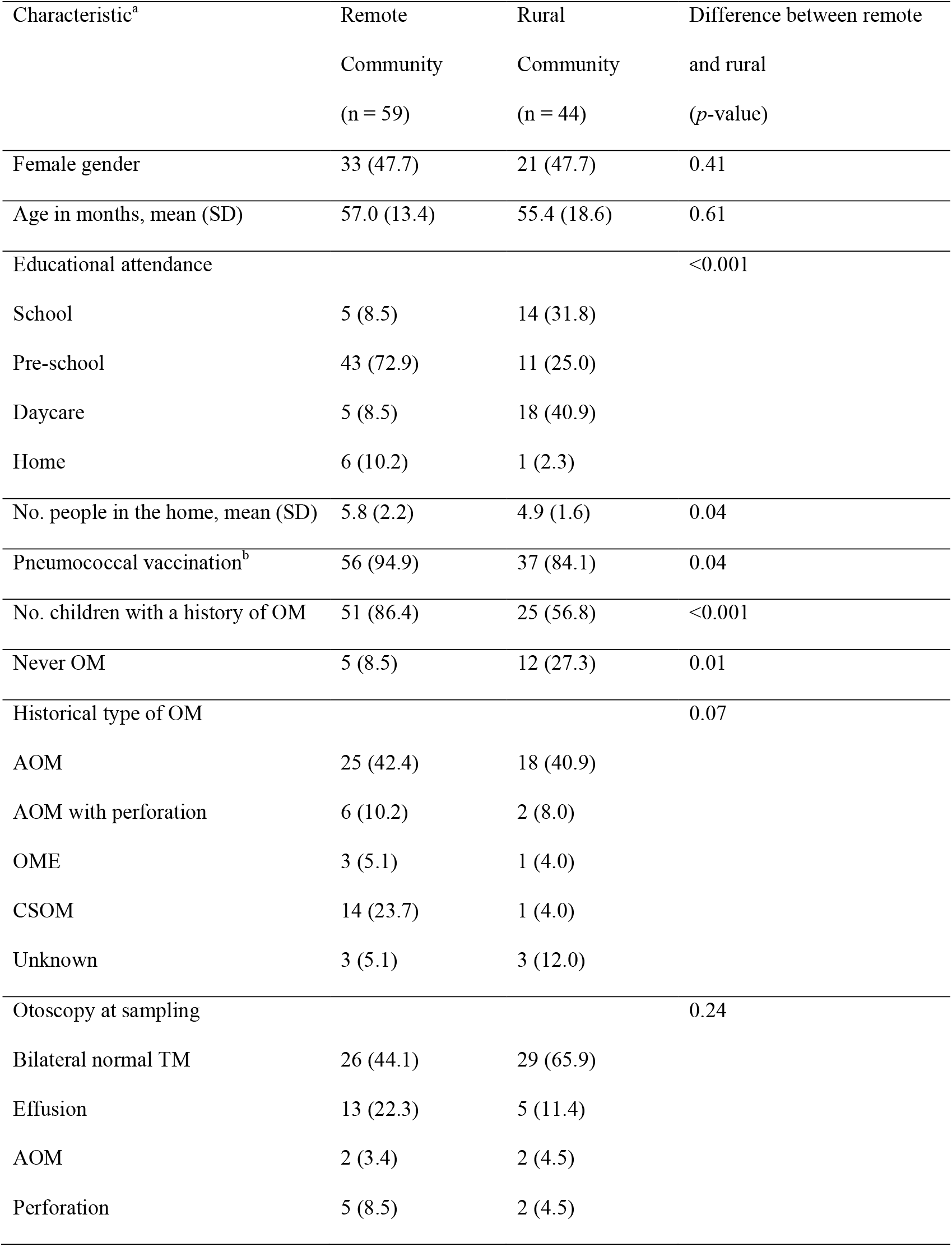
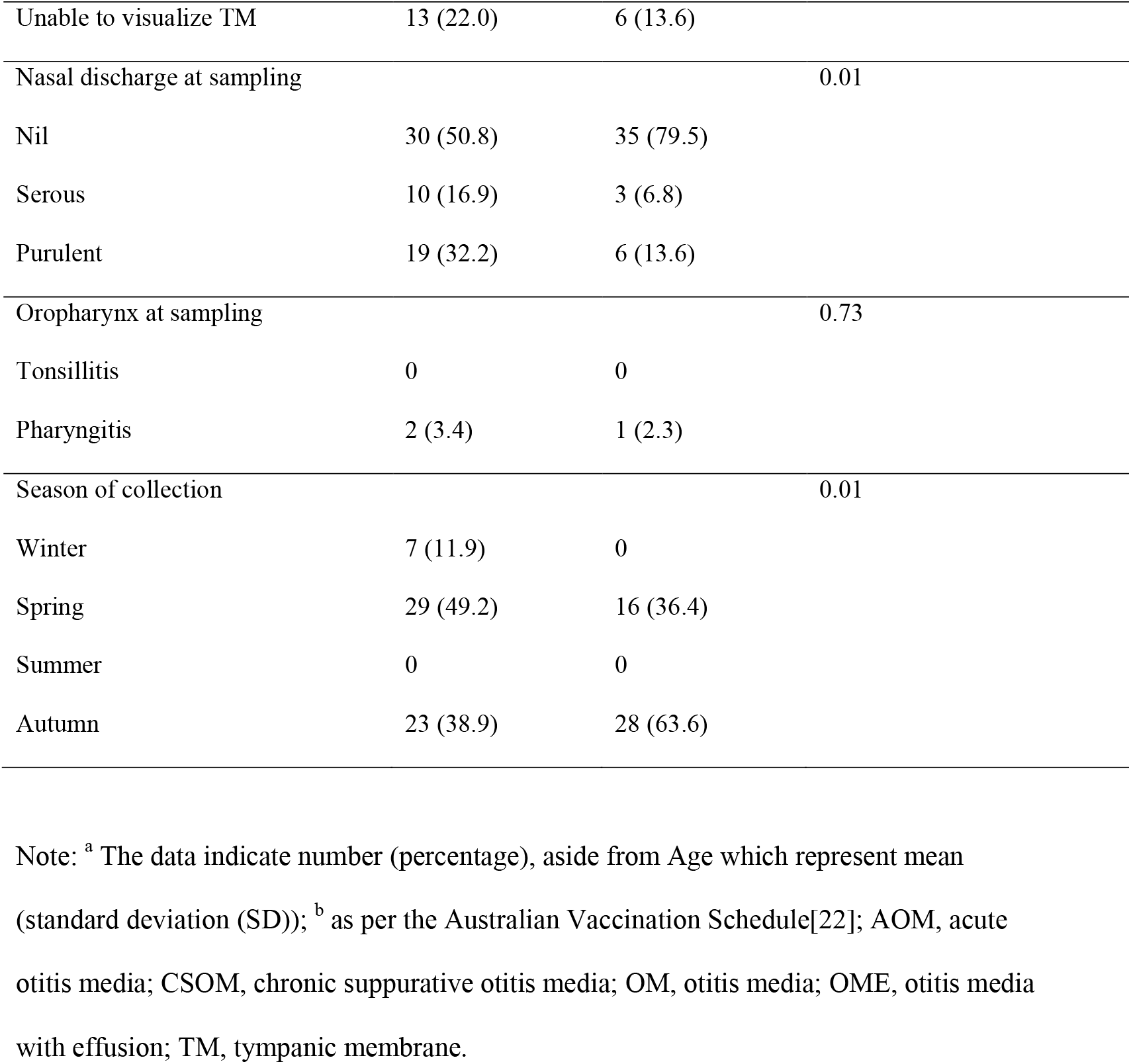
Demographic and clinical details of participants

| Characteristic<sup>a</sup> | Remote<br>Community<br>(n = 59) | Rural<br>Community<br>(n = 44) | Difference between remote<br>and rural<br>(p-value) |
| --- | --- | --- | --- |
| Female gender | 33 (47.7) | 21 (47.7) | 0.41 |
| Age in months, mean (SD) | 57.0 (13.4) | 55.4 (18.6) | 0.61 |
| Educational attendance |  |  | <0.001 |
| School | 5 (8.5) | 14 (31.8) |  |
| Pre-school | 43 (72.9) | 11 (25.0) |  |
| Daycare | 5 (8.5) | 18 (40.9) |  |
| Home | 6 (10.2) | 1 (2.3) |  |
| No. people in the home, mean (SD) | 5.8 (2.2) | 4.9 (1.6) | 0.04 |
| Pneumococcal vaccination<sup>b</sup> | 56 (94.9) | 37 (84.1) | 0.04 |
| No. children with a history of OM | 51 (86.4) | 25 (56.8) | <0.001 |
| Never OM | 5 (8.5) | 12 (27.3) | 0.01 |
| Historical type of OM |  |  | 0.07 |
| AOM | 25 (42.4) | 18 (40.9) |  |
| AOM with perforation | 6 (10.2) | 2 (8.0) |  |
| OME | 3 (5.1) | 1 (4.0) |  |
| CSOM | 14 (23.7) | 1 (4.0) |  |
| Unknown | 3 (5.1) | 3 (12.0) |  |
| Otoscopy at sampling |  |  | 0.24 |
| Bilateral normal TM | 26 (44.1) | 29 (65.9) |  |
| Effusion | 13 (22.3) | 5 (11.4) |  |
| AOM | 2 (3.4) | 2 (4.5) |  |
| Perforation | 5 (8.5) | 2 (4.5) |  |
| Unable to visualize TM | 13 (22.0) | 6 (13.6) |  |
| Nasal discharge at sampling |  |  | 0.01 |
| Nil | 30 (50.8) | 35 (79.5) |  |
| Serous | 10 (16.9) | 3 (6.8) |  |
| Purulent | 19 (32.2) | 6 (13.6) |  |
| Oropharynx at sampling |  |  | 0.73 |
| Tonsillitis | 0 | 0 |  |
| Pharyngitis | 2 (3.4) | 1 (2.3) |  |
| Season of collection |  |  | 0.01 |
| Winter | 7 (11.9) | 0 |  |
| Spring | 29 (49.2) | 16 (36.4) |  |
| Summer | 0 | 0 |  |
| Autumn | 23 (38.9) | 28 (63.6) |  |

Note: <sup>a</sup> The data indicate number (percentage), aside from Age which represent mean (standard deviation (SD)); <sup>b</sup> as per the Australian Vaccination Schedule[22]; AOM, acute otitis media; CSOM, chronic suppurative otitis media; OM, otitis media; OME, otitis media with effusion; TM, tympanic membrane.

**Figure 1:**
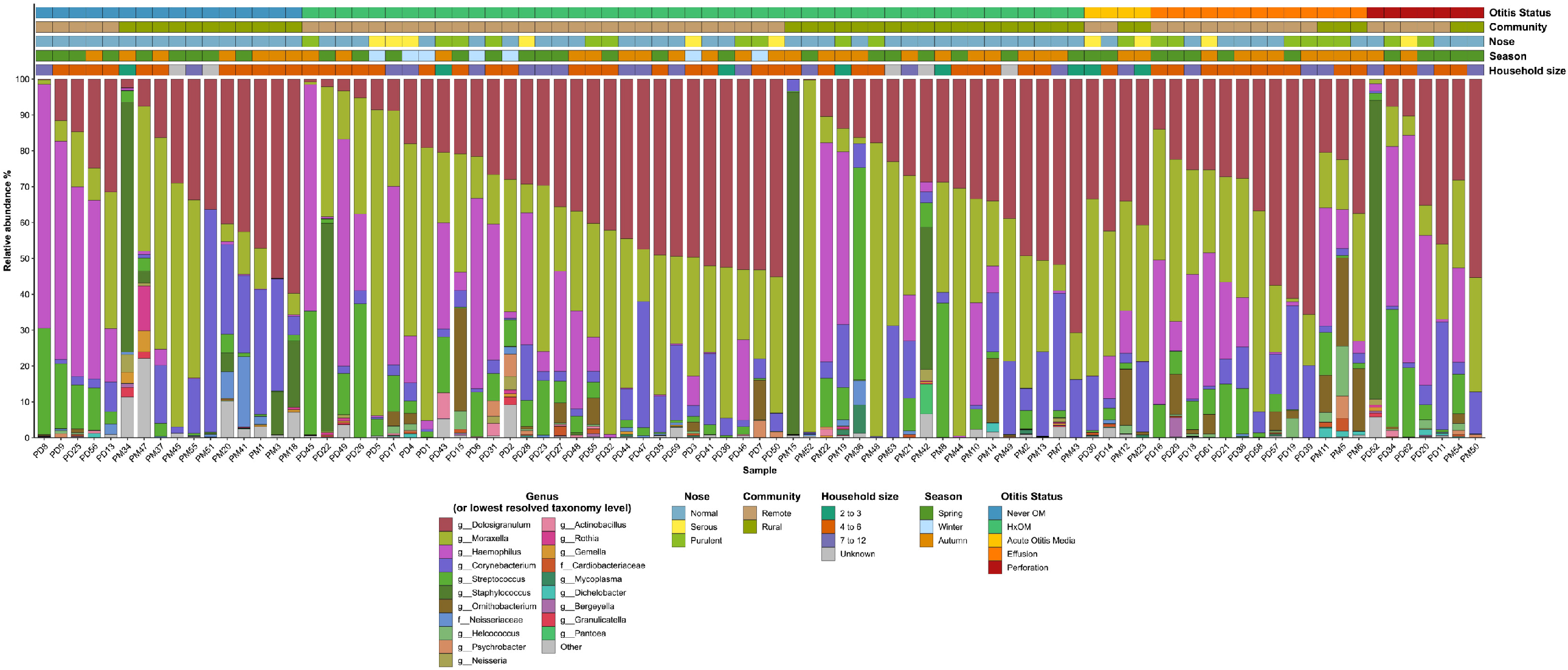
Mean relative microbial abundances of the 20 most abundant genera (or lowest resolved taxonomy level) across all samples illustrating differences between otitis Status, community of residence and other key variables. Microbes with lower abundances have been combined in the ‘Other’ (in grey). To improve interpretability, samples have been ordered by Otitis status, Community and Dolosigranulum abundance. Note: OM = otitis media, HxOM = History of OM, but health tympanic membrane at time of collection.

Network analyses showed taxa correlations largely differed according to otitis status, with the notable exception of *Streptococcus* and *Haemophilus*, which correlated across all groups. Never OM children had a more complex network of correlated genera, compared to HxOM, despite both groups having normal TMs at the time of swabbing (Figure 2). *Dolosigranulum* positively correlated with different genera across all otitis phenotypes, with exception of Effusion; to *Corynbacterium* in the Never OM group and to *Moraxella* and *Neisseriaceae* in the HxOM and TM perforation groups, respectively (Figure 2). Our culturomic data suggested the species representing the associated genera were *D. pigrum* and *C. pseudodiphtheriticum*[5]. *Ornithobacterium* correlated with *Helcococcus, Dichelobacter* (Figure 2). There were no significant differences in alpha diversity in relation to otitis status (Supplementary Table 3).

**Figure 2:**
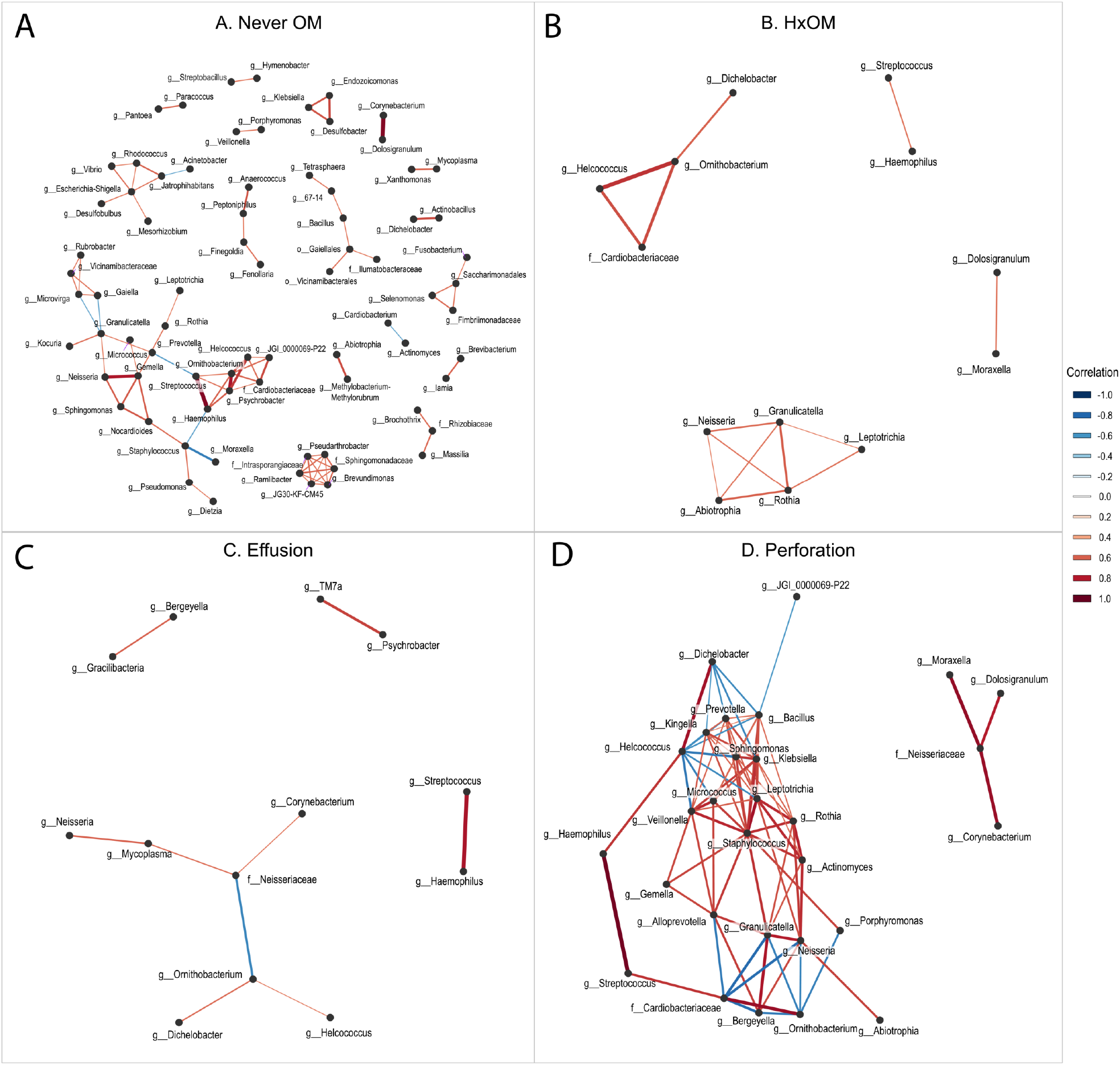
Network correlation analysis showing differences in genera relationships in context of otitis status: A) Never OM; B): HxOM; C) Middle ear effusion; D) TM perforation. Note: OM = otitis media, HxOM = History of OM, but health tympanic membrane at time of collection.

### Nasal microbiota in relation to nose health

The nasal microbiota was significantly related to nose health (PERMANOVA *p* < 0.001, F = 2.98, PERMDISP F = 2.753, *p* = 0.068). Compared to purulent rhinorrhoea, children with healthy noses had significantly higher mean relative abundance of *Staphylococcus* (6.68% vs 0.004%) and *Neisseriacea*e (0.868% vs 0.096%)(all p < 0.001) (Figure 1, Supplementary Table 2). ASV analysis showed *Staphylococcus aureus* is likely accounting for the *Staphylococcus* detections. Similar to ear health, multiple *Dolosigranulum* ASVs were detected across nose phenotypes (Supplementary Table 2). Network complexity and correlation patterns between *Dolosigranulum* and other bacteria in relation to nose status were similar to those seen within otitis status (Figure 3). *Staphylococcus* correlated negatively with *Moraxella* in healthy noses, however *Ornithobacterium* was present in all phenotypes and correlated to *Helcococcus* and *Dichelobacter* and *Cardiobacteriaceae* (Figure 3). There were no significant differences in alpha diversity in relation to nose health (Supplementary Table 3).

**Figure 3:**
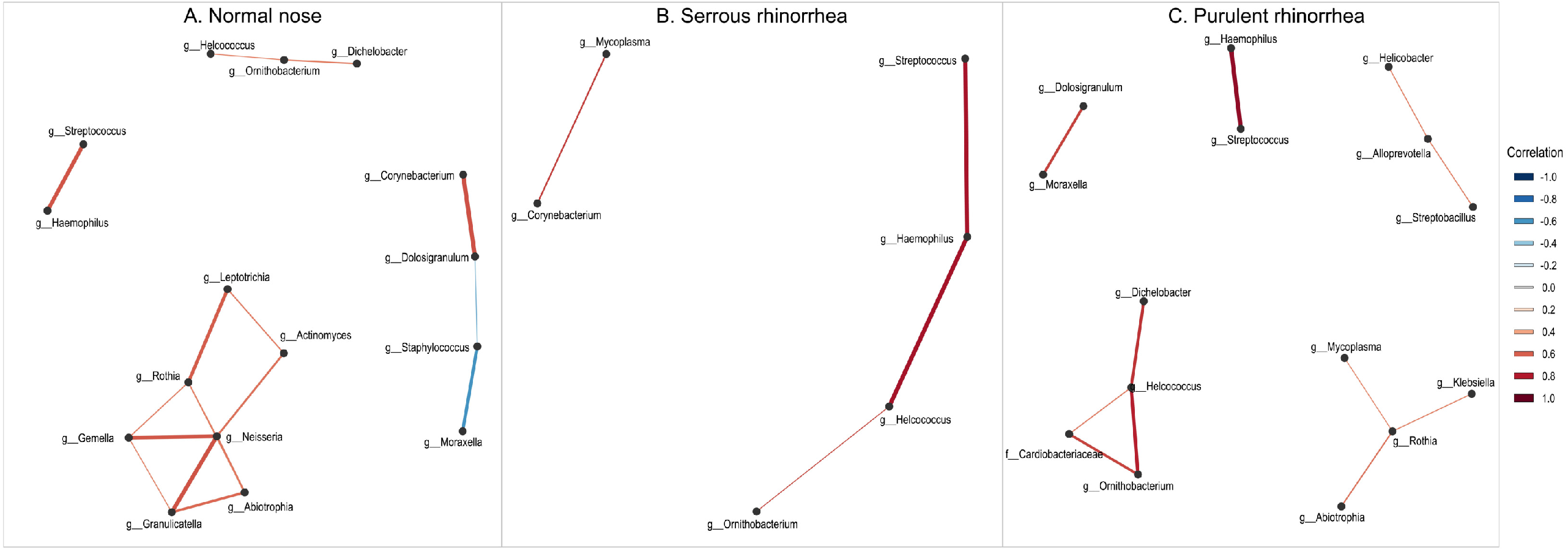
Correlation network analysis of genera in relation to nose health showing differential *Dolosigranulum* relationships.

### Nasal microbiota in relation to season, household occupancy and community

No relationship was found between nasal microbiota and season (PERMANOVA *p* = 0.456, F = 1.00; PERMDISP, *p* = 0.192, F = 1.619) or household occupancy (PERMANOVA *p* = 0.748, F = 0.791; PERMDISP *p* = 0.844, F = 0.181). There were no significant differences in relative abundance or alpha diversity for these variables (Figure 1, Supplementary Table 3). The nasal microbiota differed significantly in relation to community of residence (PERMANOVA *p* <0.001, F = 3.71), although with dispersion differences (PERMDISP, F = 7.87, *p* = 0.005). No separation was observed between the two communities on PCA (Figure 4).

**Figure 4:**
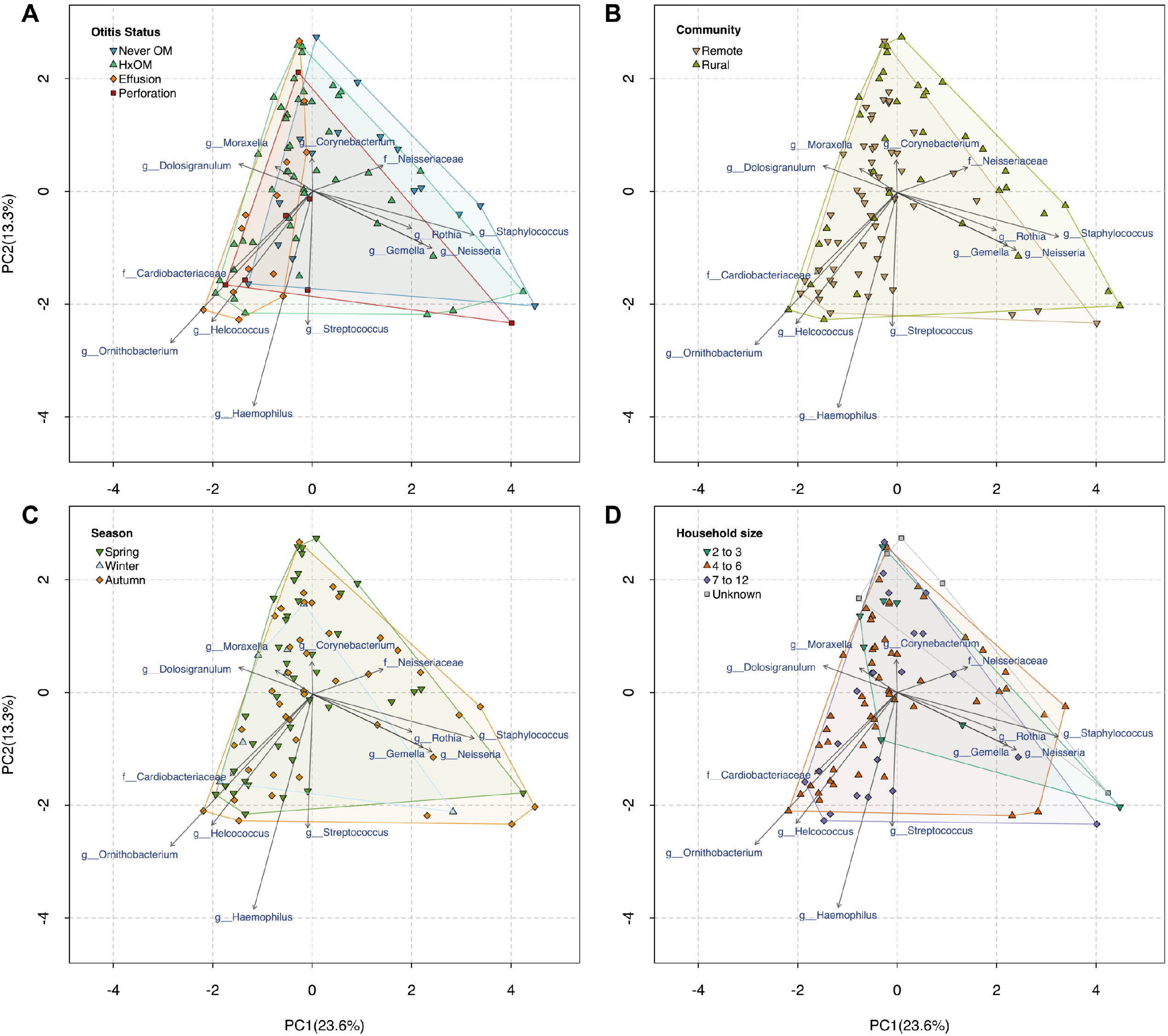
Genus-level Principal Component Analysis showing no separation in relation to A) otitis status; B) community of residence; C) season of swab collection; D) number of people residing within the household. Note: OM = otitis media, HxOM = History of OM, but healthy tympanic membrane at time of collection.

## Discussion

We demonstrated the nasal microbiota of Indigenous Australian children was related to ear and nose health. Healthy children with no history of OM showed a relationship between *Dolosigranulum* and *Corynebacterium*. We detected *Ornithobacterium* in children with OM, suggesting a potential role as a novel otopathogen in this population.

In relation to otitis status, *Moraxella* had a higher relative abundance in children with a history of OM, compared to children with no history of OM, despite both groups having healthy ears at the time of swabbing. In children with healthy noses, there was a negative correlation between *Moraxella* and *Staphylococcus*. Moraxella are common nasal colonisers whose abundance can increase during acute respiratory infections, leading to prolonged periods of enrichment within the nasal microbiota[9, 10]. Thus, the observed increase of Moraxella in HxOM children may be a downstream persistent effect of past respiratory infections (e.g. OM) leading to a remodelled microbiome distinct from children who did not contract OM. *In vitro* studies demonstrate some *Staphylococcus* species can inhibit the growth of *M. catarrhalis*[11] and may account for their negative correlation within healthy noses in our cohort.

A combination of 16S NGS and culturomic data strongly suggested there exist correlations between *C. pseudodiphtheriticum* and *D. pigrum* in healthy children with no rhinorrhoea and no historical OM. In non-Indigenous infants, *Corynebacterium* and *Dolosigranulum* are well-recognised as being associated with a stable nasopharyngeal microbiota, conferring URT and ear health[10, 12-15]. *In vitro* studies of *Corynebacterium*–*Dolosigranulum* relationships demonstrated complex interactions that were species-specific, which may be dependent on use host resources[16]. However, for the inhibition of *S. pneumoniae* both *C. pseudodiphtheriticum* and *D. pigrum* were required; neither species could inhibit the growth of *S. pneumoniae* alone[16]. These *in vitro* findings corroborate our *in vivo* data and warrant further investigation, particularly with the view towards prevention/control of otopathogen colonisation in the nose and consequent ear health benefits.

*Dolosigranulum* was ubiquitous in the nasal microbiota of our population, however examination at the level of ASV suggests this may be a heterogeneous group. The method of ASV analysis has greater precision than operational taxonomic units (OTU), used in prior URT microbiota research, and therefore may be more sensitive in detecting strain-specific differences[17]. There is one known species within the *Dolosigranulum* genus, however our findings suggest the presence of more than one strain or species, particularly given the 16S V3/V4 region as well as the wider *D. pigrum* genome has been reported to be highly conserved[16]. We did not see an inverse relationship with any of the main otopathogens, which has previously been described[18], which may be due to this population having a high baseline level of otopathogen colonisation[5]. There is growing interest in *Dolosigranulum* due to its association with URT and ear health, thus the use of whole genome sequencing and ASV analysis will provide further understanding of the nuances of this nasal commensal.

*Ornithobacterium* was absent or at low relative abundance in the nasal microbiota of children with no history of OM. This is likely to represent *O. hominis*, a newly described species of *Ornithobacterium*, which resides in the nasopharynx and is the only known human species in that genus[7]. The role of *O. hominis* in relation to respiratory/ear disease is still undetermined, however it was originally found in Australian children and Thai refugee camp infants with high rates of respiratory disease[19, 20]. Our findings suggest that *Ornithobacterium* may be associated with poor ear health. Furthermore, the network correlations supported relationships between *Ornithobacterium, Helcococcus* and *Dichelobacter* which may influence clinical outcomes. Intriguingly, the ASV and correlation network data suggests that there may be novel bacterial species within the nasal microbial ecosystem in genera which currently do not have human representatives (e.g. *Dichelobacter, Gracilibacteria*) or only have one species (*Dolosigranulum*). Along with *Ornithobacterium*, these genera warrant further investigations, particularly given their recurring relationships with genera associated with health and disease.

Although this is the largest NGS-based OM study in any indigenous population to date, limitations exist. Recruitment and sample collection in remote Australian communities is resource and time intensive, which impacted on sample size. Furthermore, recruitment of healthy children with no history of OM was challenging, despite our community-based sampling, reflecting the high burden of disease in remote Indigenous Australian communities[21]. We found that healthy children, with no history of OM, appeared to have differences in their nasal microbiota dependent on their community of residence, however the small sample size limited further sub-group analysis. A well-recognised limitation of 16S rRNA sequencing is its poor resolution at the species-level. However, a combination of ASV and culturomics data partially overcame this limitation and provided novel species-level insights into the nasal microbial ecology in nose/ear health and disease. It is hoped that moving forward methods such as metagenomic shotgun sequencing can be optimised for the URT to provide a more comprehensive assessment of the URT microbiome in relation to health and disease.

In conclusion, our investigation of the nasal microbiota of Indigenous Australian children demonstrated that there is a potential synergism between *D. pigrum* and *C. pseudodiphtheriticum* that is associated with ear and nose health. Our ASV-level analysis suggested that *Dolosigranulum* is a heterogeneous genus. Finally, we have detected the likely presence of *O. hominis* and suggestions of other novel species within the nasal microbiota that associate with poor URT/ ear health.

## Author’s contributions

ACol, AW, EGH, JA, MB, ACer, SB designed the study. ACol and AW collected samples. ACol, KC, SB conducted laboratory work. JZ and SB did the bioinformatics analysis. ACol, SB, JZ wrote the manuscript. All authors revised and approved the final manuscript.

## Transparency declaration

The authors declare that they have no conflicts of interest.

## Acknowledgments

We would like to acknowledge the valuable contributions of Jason Leon, Chantel Hunter, Gail Wason and Deborah Gertz from Mulungu Aboriginal Corporation Medical Centre, Isabel Toby from Save the Children, Doomadgee and Anne O’Keefe from Doomadgee Community Health.

